# Compensatory guaiacyl lignin biosynthesis at the expense of syringyl lignin in *4CL1*-knockout poplar

**DOI:** 10.1101/2019.12.20.885350

**Authors:** Chung-Jui Tsai, Peng Xu, Liang-Jiao Xue, Hao Hu, Batbayar Nyamdari, Radnaa Naran, Xiaohong Zhou, Geert Goeminne, Ruili Gao, Erica Gjersing, Joseph Dahlen, Sivakumar Pattathil, Michael G. Hahn, Mark F. Davis, John Ralph, Wout Boerjan, Scott A. Harding

**Affiliations:** Warnell School of Forestry and Natural Resources, University of Georgia, Athens, GA 30602, USA; Department of Genetics, University of Georgia, Athens, GA 30602, USA; Department of Plant Biology, Athens, GA 30602, USA; Center for Bioenergy Innovation, Oak Ridge National Laboratory, Oak Ridge, TN 37831, USA; Department of Plant Biotechnology and Bioinformatics, Ghent University, 9052 Ghent, Belgium; VIB Center for Plant Systems Biology, 9052 Ghent, Belgium; Department of Biochemistry, University of Wisconsin, Madison, WI 53706, USA; Great Lakes Bioenergy Research Center, Wisconsin Energy Institute, University of Wisconsin, Madison, WI 53726, USA; National Renewable Energy Laboratory, Golden, CO 80401, USA; BioEnergy Science Center, Oak Ridge National Laboratory, Oak Ridge, TN 37831, USA; Complex Carbohydrate Research Center, University of Georgia, Athens, GA 30602, USA

**Author notes:** **Author Contributions:** X.Z. generated transgenic plants and measured Klason lignin; P.X. performed histology and amplicon sequencing; J.D. measured specific gravity and acoustic velocity; H.H., S.P. and M.G.H. performed glycome profiling analysis; B.N., R.N. and S.A.H. performed glycosyl composition analysis; S.A.H. measured crystalline cellulose content; G.G. and W.B. performed phenolic profiling analysis; E.G. and M.F.D. performed saccharification analysis through the BioEnergy Science Center (BESC); R.G. and J.R. performed NMR analysis and coordinated other cell wall analysis through the Great Lakes Bioenergy Research Center (GLBRC); L.-J.X. performed bioinformatic analysis; C.-J.T. conceived the project, analyzed data and wrote the article with P.X. and S.A.H., with contributions from other authors. P.X., Department of Genetics and Informatics Institute, University of Alabama at Birmingham, Birmingham, AL 35294, USA; L-.J.X., Key Laboratory of Forest Genetics and Biotechnology, Co-Innovation Center for Sustainable Forestry in Southern China, College of Forestry, Nanjing Forestry University, Nanjing, Jiangsu 210037, China; H.H., Department of Plant Biology, Ecology, and Evolution, Oklahoma State University, Stillwater, OK 74078, USA; X.Z., State Key Laboratory of Subtropical Silviculture, Zhejiang Agriculture & Forestry University, Hangzhou, Zhejiang 310058, China; G.G., VIB Metabolomics Core, UGent-VIB Research Building, 9052 Ghent, Belgium; E.G., Los Alamos National Laboratory, Los Alamos, NM 87545, USA; S.P., Mascoma LLC (Lallemand Inc.), Lebanon, NH 03766, USA.

**Keywords:** caffeic acid, monolignol, saccharification, thiol redox, glutathione-ascorbate, sulfur assimilation, metabolic compensation

## Abstract

The lignin biosynthetic pathway is highly conserved in angiosperms, yet pathway manipulations give rise to a variety of taxon-specific outcomes. Knockout of lignin-associated *4-coumarate:CoA ligases* (*4CLs*) in herbaceous species mainly reduces guaiacyl (G) lignin and enhances cell wall saccharification. Here we show that CRISPR-knockout of *4CL1* in *Populus tremula* × *alba* preferentially reduced syringyl (S) lignin, with negligible effects on biomass recalcitrance. Concordant with reduced S-lignin was downregulation of *ferulate 5-hydroxylases* (*F5Hs*). Lignification was largely sustained by 4CL5, a low-affinity paralog of 4CL1 typically with only minor xylem expression or activity. Levels of caffeate, the preferred substrate of 4CL5, increased in line with significant upregulation of *caffeoyl shikimate esterase1*. Upregulation of *caffeoyl-CoA O-methyltransferase1* and downregulation of *F5Hs* are consistent with preferential funneling of 4CL5 products toward G-lignin biosynthesis at the expense of S-lignin. Thus, transcriptional and metabolic adaptations to *4CL1*-knockout appear to have enabled 4CL5 catalysis at a level sufficient to sustain lignification. Finally, genes involved in sulfur assimilation, the glutathione-ascorbate cycle and various antioxidant systems were upregulated in the mutants, suggesting cascading responses to perturbed thioesterification in lignin biosynthesis.

**One sentence summary:** Knockout of lignin-associated *4CL1* in *Populus* reveals a 4CL5-dependent, caffeate-modulated compensatory pathway for lignification with links to thiol redox balance and sulfur assimilation.

## INTRODUCTION

The angiosperm lignin biosynthetic pathway has been studied for decades and continues to be a topic of interest thanks to its plasticity (Sederoff et al., 1999; Boerjan et al., 2003; Ralph et al., 2004). A host of factors likely contribute to the plasticity of lignification, including enzyme specificity and taxon-dependent properties during development or in response to environmental pressures (Weng and Chapple, 2010; Vanholme et al., 2019). 4-Coumarate:CoA ligase (4CL) catalyzes ATP-dependent CoA-thioesterification of various cinnamic acid derivatives (Knobloch and Hahlbrock, 1975) in arguably the most promiscuous step of monolignol biosynthesis. In all sequenced angiosperm genomes, 4CL is encoded by multiple genes belonging to two distinct phylogenetic classes (Ehlting et al., 1999; Saballos et al., 2012; Chen et al., 2014). Although Class II 4CLs involved in the biosynthesis of flavonoids and other soluble phenolics exhibit a substrate preference for 4-coumaric acid, many lignin-associated Class I members show similar *in vitro* affinities for caffeic acid, 4-coumaric acid, and sometimes ferulic acid (Ehlting et al., 1999; Harding et al., 2002; Lindermayr et al., 2002; Gui et al., 2011; Chen et al., 2013).

The *Populus* genome contains four Class I *4CL* members as two paralogous pairs derived from Salicoid whole-genome duplication (Tsai et al., 2006; see Supplemental Figure 1). Among them, *4CL1* (Potri.001G036900) is known to be involved in lignin biosynthesis based on molecular and reverse genetic characterization in *P. tremuloides* Michx. and *P. tremula* × *alba* INRA 717-1B4 (Hu et al., 1998; Hu et al., 1999; Voelker et al., 2010). The genome duplicate *4CL5* (Potri.003G188500) is the only other *4CL* gene family member expressed in lignifying xylem, but its transcript levels are much lower than those of *4CL1* (Supplemental Figure 1; Hefer et al., 2015; Swamy et al., 2015; Hu et al., 2016; Xue et al., 2016; Wang et al., 2018). *Populus trichocarpa* (Nisqually-1) *4CL1* and *4CL5* were proposed to encode 4CL proteins that form heterotetramers in a 3:1 ratio (referred to as Ptr4CL3 and Ptr4CL5 in Chen et al., 2013; Chen et al., 2014). The activity of individual isoforms as well as the tetrameric complex is sensitive to inhibition by hydroxycinnamic acids and their shikimate esters (Harding et al., 2002; Chen et al., 2014; Lin et al., 2015). Such complexity of 4CL catalysis is consistent with multiple cellular strategies for directing hydroxycinnamic acids toward lignin biosynthesis. Furthermore, silencing lignin-associated *4CLs* can have different effects on syringyl-to-guaiacyl lignin (S/G) ratio depending on the species, ranging from increases in tobacco, *Arabidopsis*, and switchgrass (Kajita et al., 1996; Lee et al., 1997; Xu et al., 2011), to little change in alfalfa (Nakashima et al., 2008), to decreases in and rice (Gui et al., 2011). Variable effects on S/G ratio have even been reported among closely related poplar species (Hu et al., 1999; Voelker et al., 2010; Chanoca et al., 2019 and references therein). Such variation has been attributed both to differing degrees of 4CL downregulation, as well as to 4CL multiplicity (Boerjan et al., 2003; Saballos et al., 2012). Analysis of knockout (KO) mutants should eliminate the uncertainty caused by partial gene silencing.

**Figure 1.**
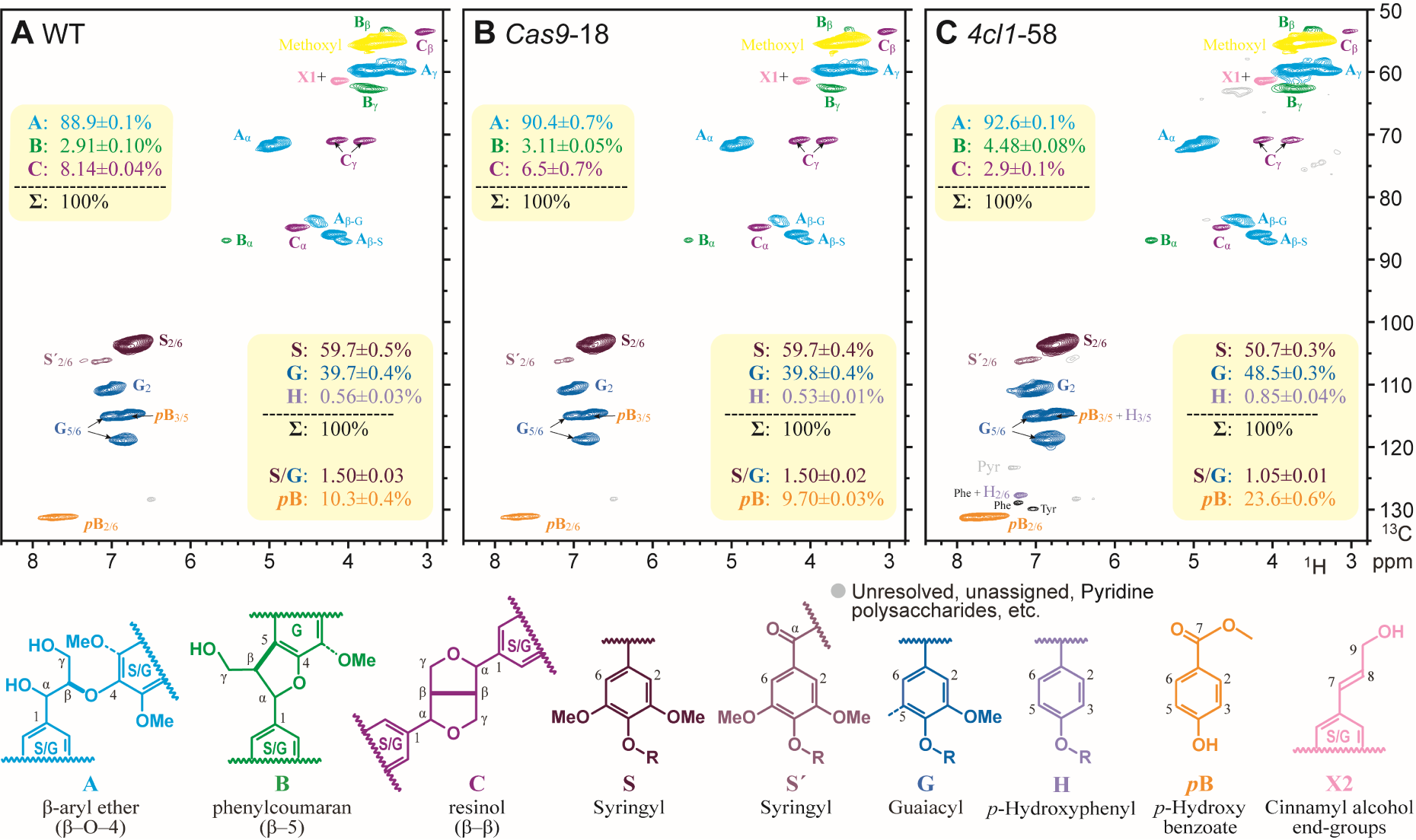
NMR analysis of *4cl1* mutant and control poplar wood. Representative ^1^H–^13^C (HSQC) correlation spectra of the aromatic region of enzyme lignins from ball-milled control (WT, left, or Cas9, middle) and a *4cl1* mutant (right) wood samples. The main lignin structures and linkages identified are illustrated below and color-coded to match their assignments in the spectra. Volume integrals (with the same color coding) were measured using the α-C/H correlation peaks from **A, B**, and **C** units, and **S**_2/6_ + **S’**_2/6_, **G**2, and **H**_2/6_ (corrected for Phe) aromatics (with the integrals halved as usual for the **S, H**, and **C** units) are noted as the mean ± SE of biological replicates (n=3 for WT, 2 for *Cas9*, and 5 for *4cl1*) (Kim and Ralph, 2010; Mansfield et al., 2012). Note that the interunit linkage distribution (**A**:**B**:**C**) is determined from volume integrals of just those units and made to total 100%; small differences (such as the apparently enhanced β-ether **A** level in the *4cl1* line) should not be overinterpreted, as we were unable to delineate authenticated peaks for, nor therefore obtain reliable accounting of, various tetrahydrofurans (from β–β-coupling of monolignol *p*-hydroxybenzoates), nor of the 4–O–5- and 5–5-linked units that require one or two G units (and would therefore logically be higher in the *4cl1* lines, with their higher %G-units). Also note that the H-unit (H_2/6_) correlation peak overlaps with another peak from Phe protein units (Kim et al., 2017); integrals were corrected by subtracting the integral from the resolved Phe peak below it to obtain the best estimate available.

*Arabidopsis*, sorghum, and maize *4cl* mutants showed preferential reductions in G lignin, resulting in increased S/G ratios (Saballos et al., 2012; Van Acker et al., 2013; Li et al., 2015; Xiong et al., 2019). Though rare until recently, transgenic nulls can now be efficiently obtained for genetically less tractable systems like woody perennials or polyploids using CRISPR/Cas9 technology (Voytas and Gao, 2014; Bewg et al., 2018). We previously reported that CRISPR-KO of the predominant lignin *4CL* in poplar led to a reduced S/G ratio (Zhou et al., 2015), whereas similar KO in tetraploid switchgrass increased the S/G ratio (Xu et al., 2011). The present study aims to further characterize the poplar *4cl1* mutants by more comprehensive cell wall analysis, biomass saccharification, phenolic profiling, and RNA-Seq. We present data to show distinct effects of *4CL-*KO on lignin biosynthesis and enzymatic hydrolysis of cell walls relative to *4cl* mutants of *Arabidopsis* and other herbaceous species. Our findings suggest a high sensitivity of caffeic acid homeostasis to 4CL1 perturbation which enables 4CL5 catalysis and compensatory production of G-enriched lignin in the *Populus* mutants.

## RESULTS

### *4CL1-*knockout preferentially reduces S lignin biosynthesis

We reported previously that CRISPR editing of *4CL1* in *Populus tremula* x *alba* INRA 717-1B4 (hereafter *4cl1* mutants) led to uniformly discolored wood and a 23% reduction in lignin content based on pyrolysis molecular beam mass spectrometry (pyMBMS) analysis (Zhou et al., 2015). Total lignin content determined by the Klason method revealed a 19% reduction in transgenic wood (Table 1). The pyMBMS analysis also detected a 30% decrease in the S/G ratio (Zhou et al., 2015), attributable to a sharp (34%) reduction of S units, and a small (5%) though significant reduction of G units (Table 1). Thioacidolysis showed a similar response in S lignin, but a greater reduction (18%) of G lignin, in addition to a 52% increase of the minor component, H lignin (Table 1), which is poorly pyrolyzed and hence undetectable using pyMBMS (Sykes et al., 2015). To probe the structural modifications in walls of the mutant further, 2D-NMR analysis was performed using both whole cell wall residues and cellulolytic enzyme lignins. The results were highly concordant with those of pyMBMS and thioacidolysis, showing a ∼30% decrease in S/G ratio along with increased H lignin in the mutants (Figure 1). The frequency of *p*-hydroxybenzoate units also increased in the mutants (Figure 1), but we do not see evidence for the appearance of (higher levels of) β–β-coupling products derived from monolignol (largely sinapyl) *p*-hydroxybenzoate conjugates to produce tetrahydrofurans rather than the familiar resinols **C** derived from unconjugated monolignols, as has been observed in palms (Lu et al., 2015). Taken together, all analytical methods employed in this study showed consistent decreases of S/G ratio by 20-30% in the *4cl1* mutants, resulting from a preferential reduction of S lignin.

**Table 1.**
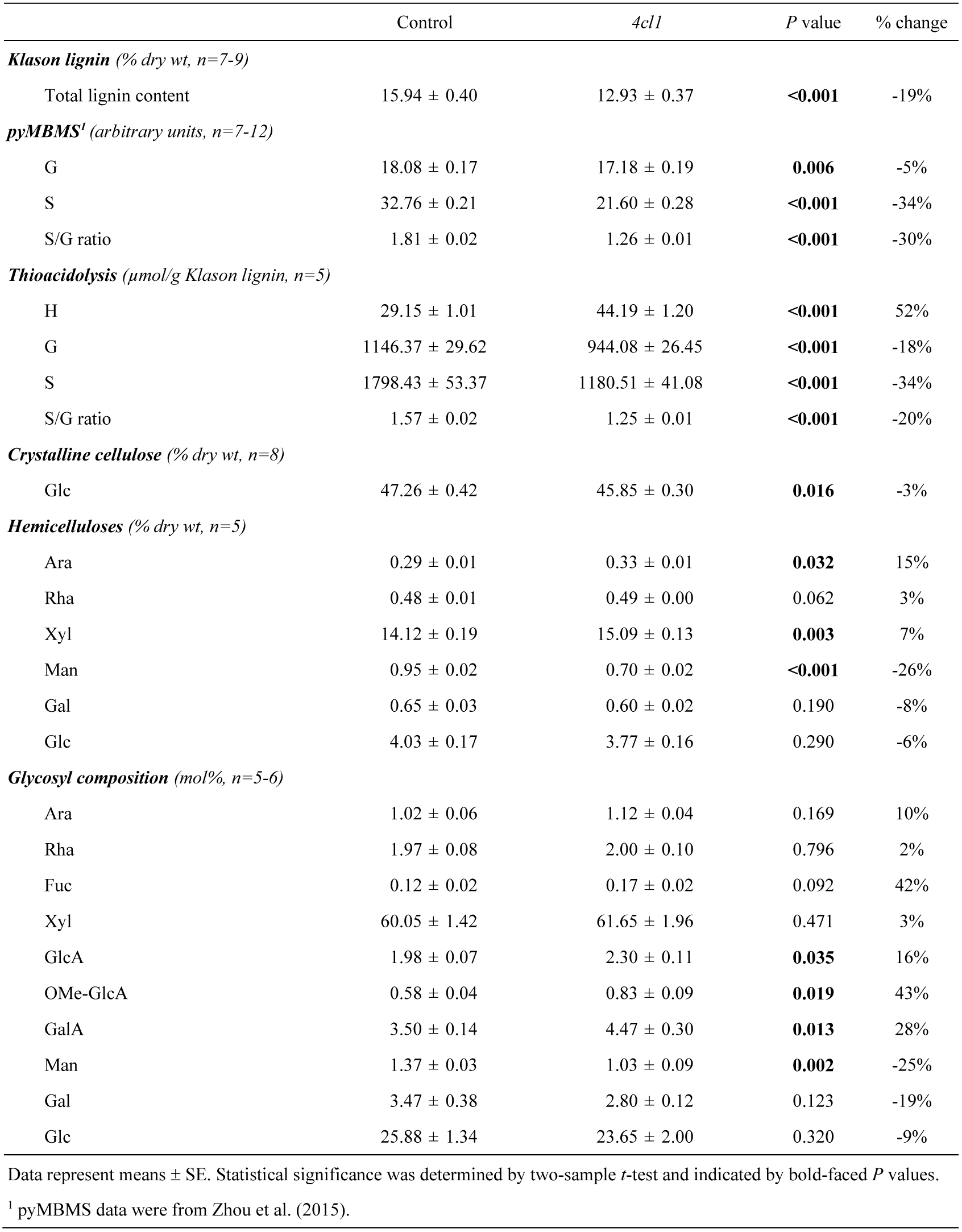
Wood chemical properties of control and *4cl1*-KO poplars.

Phloroglucinol staining of stem cross sections confirmed reduced lignification in the mutants (Figures 2A and 2B). In agreement with previous findings (Coleman et al., 2008; Voelker et al., 2010), reduced lignin accrual led to collapsed xylem vessels (Figures 2B and 2D). Accordingly, wood specific gravity was significantly reduced in *4cl1* mutants (Figure 2E). The mutants also exhibited significantly lower acoustic velocity (Figure 2F), which is correlated with the microfibril angle of the S2 layer (Schimleck et al., 2019). No other apparent growth anomaly was observed under greenhouse conditions.

**Figure 2.**
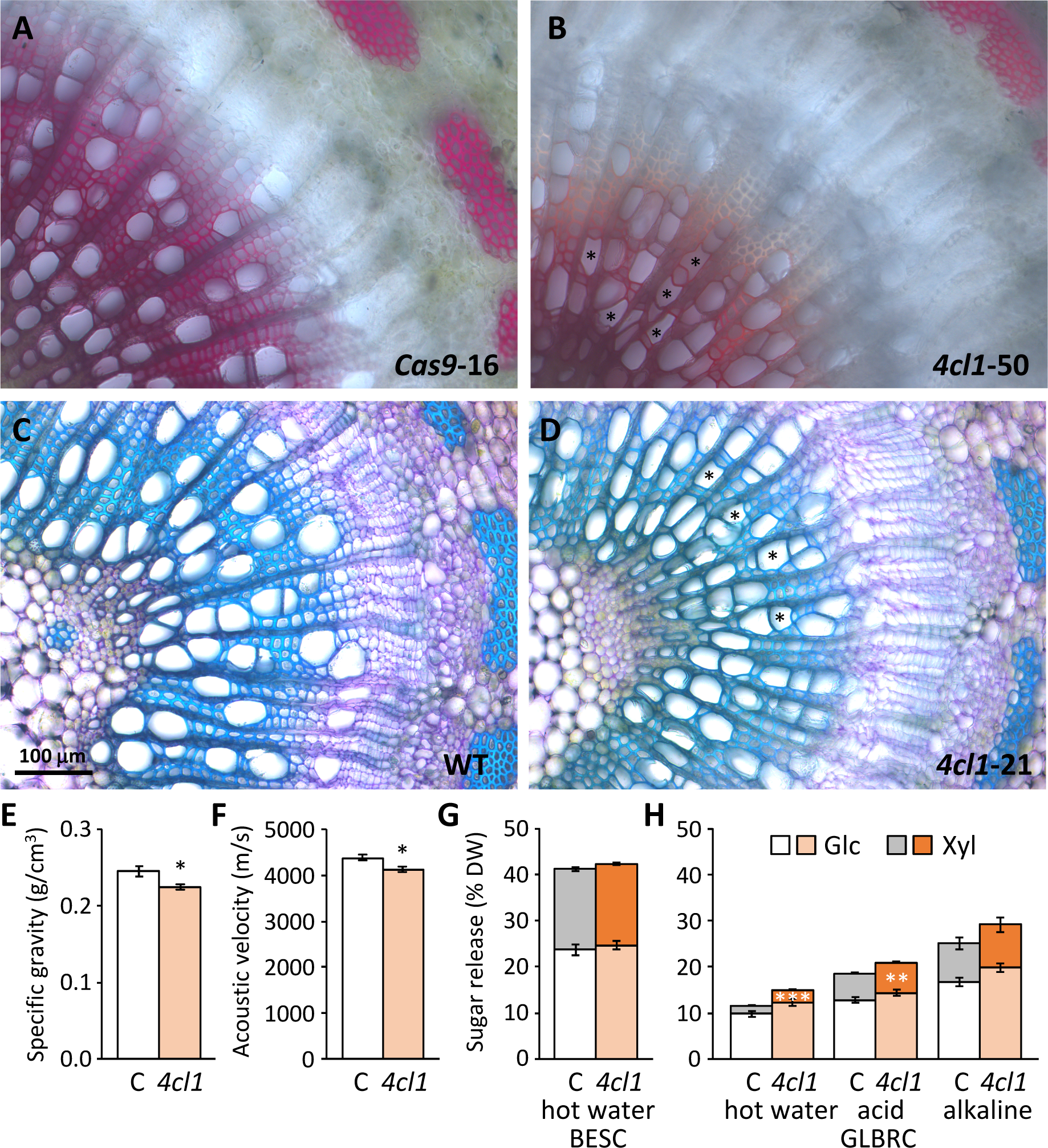
Histology, physical properties and saccharification of mutant and control poplar wood. **A-B**, Stem cross sections (10^th^ internode) stained with phloroglucinol. **C-D**, Stem cross sections (8^th^ internode) stained with toluidine blue. Representative images from control (WT or Cas9) and *4cl1* mutant lines are shown. Collapsed vessels are marked with asterisks. Scale bar = 100 μm. **E-F**, Wood specific gravity (**E**) and acoustic velocity (**F**) of control (n=9) and *4cl1* (n=12) samples. **G-H**, Enzymatic sugar release of control and mutant samples following BESC hydrothermal (**G**, n=6) and GLBRC hydrothermal, dilute acid and dilute alkaline pretreatments (**H**, n=5). Data in E-H represent means±SE. Statistical significance was determined by Student’s *t* test (***, *P* <0.001; **, *P* <0.01; *, *P* <0.05).

### *4CL1-*knockout has no obvious effects on biomass saccharification

Reduction of lignin content, regardless of S/G change, has been shown to improve enzymatic sugar release in alfalfa, *Arabidopsis*, and poplar (Chen and Dixon, 2007; Mansfield et al., 2012; Van Acker et al., 2013; Wang et al., 2018). Unexpectedly, we found no difference between control poplar and *4cl1* mutants in enzymatic glucose and xylose release following hydrothermal pretreatment (Figure 2G) using the high-throughput recalcitrance assay developed by the US Department of Energy (DOE)-funded BioEnergy Science Center (BESC) (Selig et al., 2010). Independent analyses using the high-throughput method of the DOE-Great Lakes Bioenergy Research Center (GLBRC) (Santoro et al., 2010) also detected no differences in glucose release (Figure 2H). Mutant samples released slightly higher levels of xylose, but only after less severe (hot water and dilute acid, not alkaline) pretreatments (Figure 2H). The overall lower sugar yields from the GLBRC assays are similar to those reported elsewhere (Wilkerson et al., 2014), and reflect the milder pretreatment and enzymatic hydrolysis conditions than those applied with the BESC methods (Santoro et al., 2010; Selig et al., 2010).

### Altered lignin-carbohydrate interactions in the cell wall of *4cl1* mutants

As lignin is purportedly cross-linked with cell wall polysaccharides, we next examined the *4CL1*-KO effects on cell wall glycans and their interaction with lignin. We detected a small decrease in crystalline cellulose contents of the mutants (Table 1). Monosaccharides typically associated with hemicelluloses showed slight changes, with mannose decreased significantly, and the most abundant xylose increased slightly (Table 1). Glycosyl residue composition analysis further revealed increased mol% of glucuronic acid (GlcA), its 4-*O*-methylated form (OMe-GlcA), and galacturonic acid (GalA), the main component of pectins (Table 1). These data suggest altered cell wall polysaccharide composition in lignin-reduced *4cl1* mutants.

Extractive-free wood meals were then subjected to cell wall fractionation to provide a finer resolution analysis of matrix polysaccharide organization by glycome profiling using a panel of cell wall glycan-directed monoclonal antibodies (Pattathil et al., 2010; Pattathil et al., 2012). The amounts of extractable cell wall material differed little between control and *4cl1* mutants (Supplemental Figure 2A). The glycan epitope profiles from cell wall fractions enriched with pectins (oxalate and carbonate extracts) and hemicelluloses (1 M and 4 M KOH extracts) were largely similar between genotypes, with a few epitopes exhibiting slightly lower signals in the mutants (Figure 3, Supplemental Figure 2B). In contrast, we observed significantly increased polysaccharide extractability in the chlorite fraction of mutant cell walls (Figure 3), especially for epitopes recognized by mAb groups Xylan-5 (4-*O*-methyl GlcA-substituted xylans), Xylan-6/Xylan-7 (linear unsubstituted xylans), RG-I (rhamnogalacturonan I), RG-I/AG (arabinogalactan), and pectic (RG-I and homogalacturonan) backbones (Pattathil et al., 2010; Schmidt et al., 2015; Ruprecht et al., 2017). Because chlorite degrades lignin and liberates lignin-bound glycans, the enhanced carbohydrate extractability in the chlorite fraction is consistent with less interaction between matrix polysaccharides and lignin in the mutant cell wall. Whether the increased release of those glycans reflects reduced lignin abundance and/or altered lignin composition was not determined.

**Figure 3.**
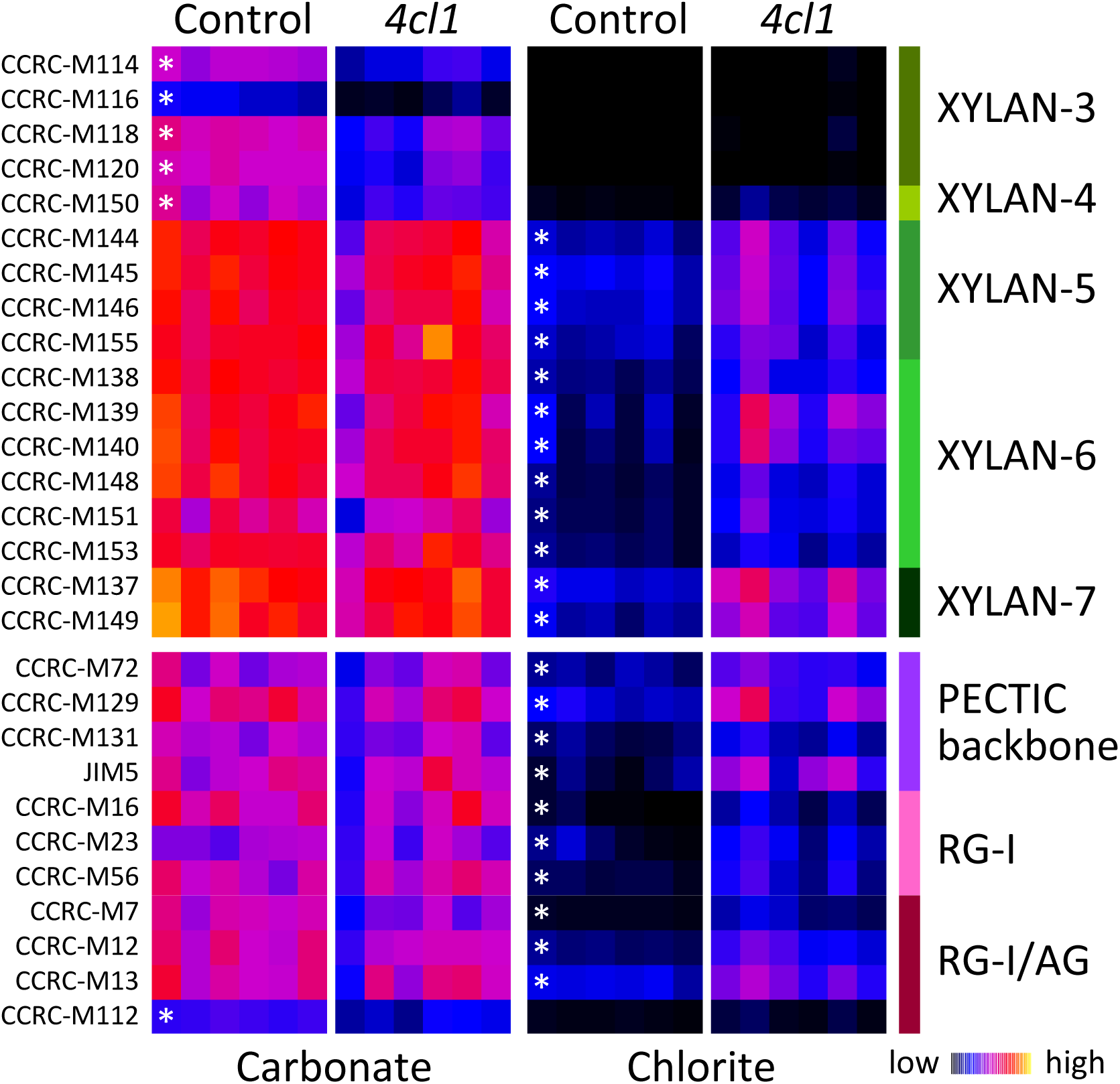
Glycome profiling of *4cl1* mutant and control poplar wood. Heatmap depiction of signal intensities resulting from binding of cell wall glycan-directed mAbs to two cell wall fractions extracted by sodium carbonate and chlorite from control and mutant samples. mAbs are arranged in rows by the cell wall glycan epitope groups (vertical bars on the right), and biological replicates (n=6) are shown in columns. Asterisks indicate significant differences between plant groups determined by Student’s *t* test (*P* <0.01, fold-change ≥1.5).

### Altered phenylpropanoid metabolism in *4cl1* mutants

UPLC-MS profiling of methanolic extracts from developing xylem revealed a major shift in phenylpropanoid metabolism in the mutants. Consistent with decreased lignin biosynthesis, various oligolignols and their hexosides were detected at drastically reduced levels in the mutants (Table 2). Also significantly reduced were lignin pathway intermediates 4-coumaroyl and caffeoyl shikimate esters (Table 2), whose synthesis depends on 4CL-activated hydroxycinnamoyl-CoAs as acyl donors (Hoffmann et al., 2003). Among potential 4CL substrates, caffeic acid is the only hydroxycinnamic acid consistently detected in control xylem under our experimental conditions (Table 2). The *4cl1* mutants accrued elevated levels of caffeic acid and its sulfate ester, along with very high levels of phenolic hexose esters and hexosides (Table 2). In particular, a caffeic acid 3/4-*O*-hexoside increased by nearly 80-fold and became the most abundant soluble phenylpropanoid identified in the mutant xylem extracts (Table 2). Mutants also contained detectable levels of 4-coumaric acid and ferulic acid sulfate, as well as highly increased hexose conjugates of 4-coumarate, ferulate and sinapate (Table 2). Chlorogenic acid (5-*O*-caffeoyl quinate), the predominant hydroxycinnamoyl quinate ester in poplar, changed little in the mutants, whereas the less abundant 3-*O*-caffeoyl quinate, 3-*O*-feruloyl quinate and 4-*O*-feruloyl quinate increased significantly (Table 2). It appears that, unlike for the shikimate conjugates, synthesis of hydroxycinnamoyl quinate esters in poplar is largely independent of 4CL1. Together, the phenolic profiling data revealed a buildup of hydroxycinnamates and their diversion into nonstructural phenylpropanoid pools at the expense of lignin precursors in the *4cl1* mutants.

**Table 2.** Xylem phenolic metabolites with significantly altered abundance in *4cl1* mutants.

| Name | RT | m/z | ion | Control | *4cl1* | Fold change |
| --- | --- | --- | --- | --- | --- | --- |
| ***oligolignol*** |  |  |  |  |  |  |
| G(8-*O*-4)G 9-*O*-hexoside | 7.15 | 537.1995 | M-H | 1820 ± 123 | 47 ± 22 | -39 |
| G(8-*O*-4)G hexoside | 7.44 | 537.1995 | M-H | 7153 ± 430 | 1406 ± 167 | -5.1 |
| G(8-*O*-4)G(8-*O*-4)G | 9.94 | 571.2202 | M-H | 3242 ± 190 | 3 ± 2 | -1064 |
| Gox(8-*O*-4)G | 11.92 | 373.1307 | M-H | 1321 ± 70 | 12 ± 4 | -110 |
| G(8-5)G [-H<sub>2</sub>O] | 13.24 | 339.1250 | M-H | 2049 ± 181 | 23 ± 10 | -89 |
| G(8-*O*-4)S(8-5)G | 14.95 | 583.2209 | M-H | 6137 ± 402 | 65 ± 49 | -94 |
| G(8-*O*-4)S(8-5)G | 15.77 | 583.2205 | M-H | 1264 ± 154 | 2 ± 1 | -641 |
| ***hydroxycinnamoyl shikimate*** |  |  |  |  |  |  |
| 4-coumaroyl shikimate | 9.72 | 319.0841 | M-H | 17617 ± 4276 | 4658 ± 734 | -3.8 |
| caffeoyl shikimate | 6.28 | 335.0822 | M-H | 79601 ± 16211 | 16973 ± 1928 | -4.7 |
| caffeoyl shikimate | 7.78 | 335.0791 | M-H | 16415 ± 2815 | 2844 ± 324 | -5.8 |
| feruloyl shikimate | 10.31 | 349.0942 | M-H | 682 ± 140 | 23 ± 10 | -30 |
| ***phenolic acid*** |  |  |  |  |  |  |
| 4-coumaric acid<sup>^</sup> | 7.96 | 163.0399 | M-H | 41 ± 13 | 760 ± 114 | 19 |
| caffeic acid | 5.04 | 179.0352 | M-H | 1110 ± 121 | 3404 ± 389 | 3.1 |
| caffeic acid sulfate | 1.82 | 258.9947 | M-H | 1392 ± 138 | 4650 ± 254 | 3.3 |
| ferulic acid sulfate | 5.07 | 273.0085 | M-H | 0 ± 0 | 3912 ± 315 | ∞ |
| ***phenolic acid hexose/hexoside*** |  |  |  |  |  |  |
| 4-coumaroyl hexose | 4.64 | 325.0996 | M-H | 2876 ± 431 | 212766 ± 15741 | 74 |
| 4-coumaroyl hexose | 5.21 | 325.0985 | M-H | 1994 ± 334 | 88055 ± 5699 | 44 |
| caffeoyl hexose | 3.43 | 341.0931 | M-H | 224 ± 103 | 13415 ± 936 | 60 |
| caffeic acid 3/4-*O*-hexoside | 4.13 | 341.0946 | M-H | 7950 ± 649 | 611832 ± 22149 | 77 |
| caffeic acid 3/4-*O*-hexoside | 5.61 | 341.0937 | M-H | 6539 ± 302 | 205906 ± 12360 | 32 |
| ferulic acid 4-*O*-hexoside | 3.83 | 711.2309 | 2M-H | 117 ± 56 | 209459 ± 10490 | 1798 |
| feruloyl hexose | 5.46 | 355.1104 | M-H | 1539 ± 415 | 184710 ± 11173 | 120 |
| feruloyl hexose | 5.54 | 711.2215 | 2M-H | 0 ± 0 | 32748 ± 3070 | ∞ |
| feruloyl hexose | 5.96 | 355.1093 | M-H | 266 ± 144 | 38777 ± 3127 | 146 |
| sinapic acid 4-*O*-hexoside | 4.49 | 771.2552 | 2M-H | 1463 ± 217 | 263304 ± 8064 | 180 |
| sinapic acid 4-*O*-hexoside | 6.21 | 385.1179 | M-H | 1091 ± 89 | 78135 ± 5624 | 72 |
| 4-hydroxybenzoyl hexose | 2.74 | 299.0824 | M-H | 4356 ± 426 | 95399 ± 9034 | 22 |
| vanillic acid 4-*O*-hexoside | 2.30 | 659.1967 | 2M-H | 9322 ± 313 | 363392 ± 13301 | 39 |
| ***hydroxycinnamoyl quinate*** |  |  |  |  |  |  |
| 3-*O*-caffeoyl quinate | 3.11 | 353.0889 | M-H | 2001 ± 284 | 7236 ± 833 | 3.6 |
| 3-*O*-feruloyl quinate | 4.83 | 367.1046 | M-H | 1026 ± 127 | 2665 ± 341 | 2.6 |
| 4-*O*-feruloyl quinate | 6.83 | 367.1047 | M-H | 393 ± 85 | 1746 ± 194 | 4.4 |
| 5-*O*-caffeoyl quinate<sup>§</sup> | 4.32 | 353.0911 | M-H | 21688 ± 2000 | 28126 ± 2621 | 1.3 |
| 4-*O*-caffeoyl quinate<sup>§</sup> | 4.69 | 353.0893 | M-H | 2283 ± 220 | 2630 ± 266 | 1.2 |

Data represent mean signal intensities ± SE of n= 10 control (3 WT and 7 Cas9 vector control lines) or 14 *4cl1*-KO lines. Only data with signal intensities ≥1000 in at least one group (except <sup>^</sup>), significant differences (*P* <0.002) between groups based on Student's *t*-test (except <sup>§</sup>), and confirmed peak annotation are shown. The exceptions are included for reference.

### Transcriptional adjustments of lignin biosynthesis

RNA-Seq analysis identified 1752 genes with significantly altered xylem transcript abundance in response to *4CL1*-KO; about two-thirds of which were upregulated (1153) and the remaining (599) downregulated. Because guide RNA (gRNA)-directed CRISPR/Cas9 cleavages and frameshift mutations are located in the first exon of Pta*4CL1* (Zhou et al., 2015), aberrant transcripts containing premature stop codons would be expected in the mutants and likely be subject to nonsense-mediated mRNA decay (Conti and Izaurralde, 2005). Accordingly, *4CL1* was among the most significantly downregulated genes in the mutants, with its transcript levels decreased by 92% (Figure 4A). Our data thus underscore the efficacy of CRISPR/Cas9 mutagenesis at multiple levels.

**Figure 4.**
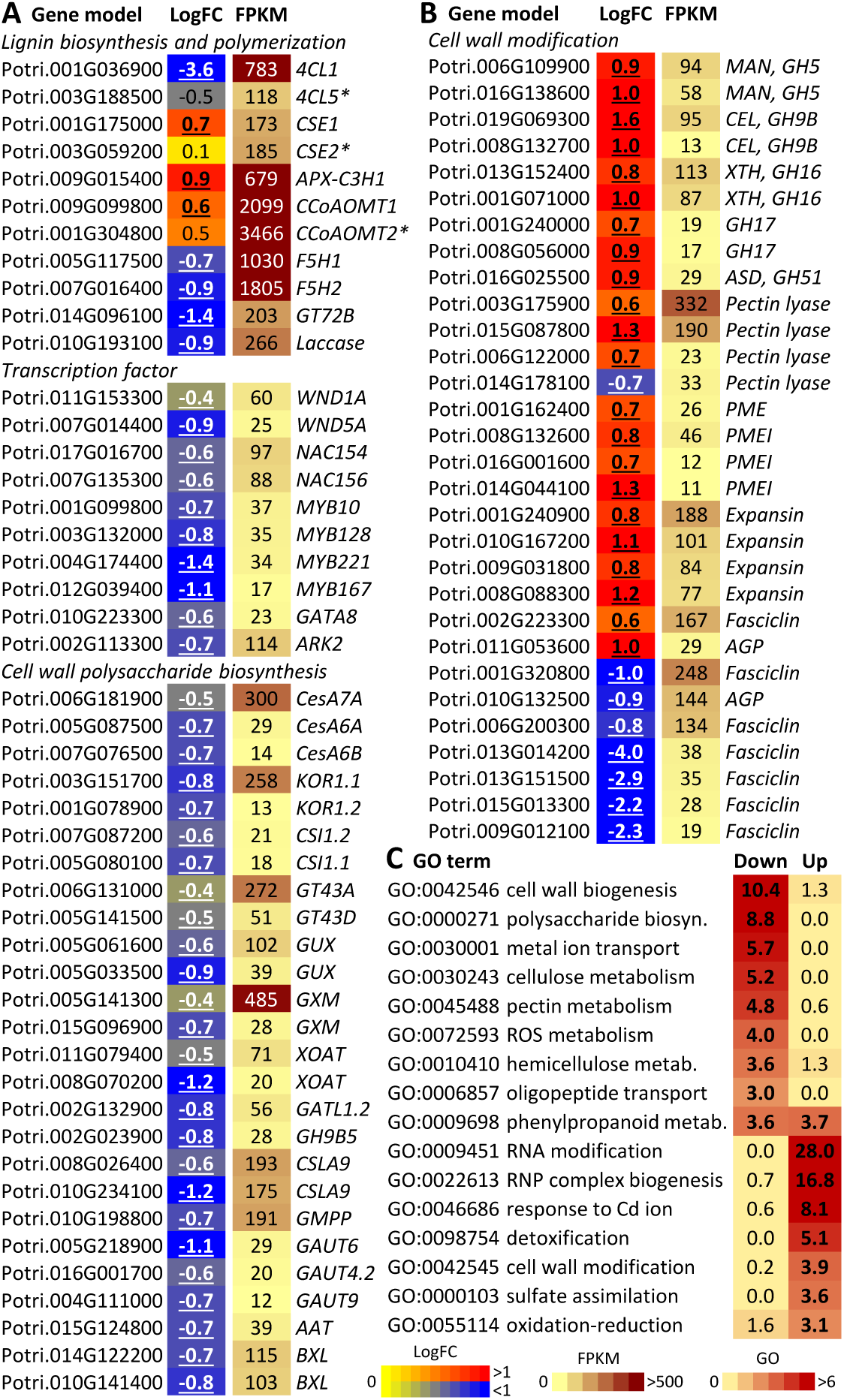
Transcriptional responses of *4cl1* mutants. **A**, Expression response heatmaps of genes involved in cell wall biogenesis. **B**, Expression response heatmaps of genes involved in cell wall remodeling. LogFC, log_2_-transformed fold-change (mutant/control) values; FPKM, average transcript abundances of control samples. Only genes with significant differences (*Q* ≤0.01, boldface and underlined) and with control FPKM ≥10 are shown, except for genome duplicates marked with asterisks. **C**, GO enrichment of differentially up- or down-regulated genes in the mutants. Representative GO terms are shown with the negative log_10_-transformed *P* values. Heatmaps are visualized according to the color scales at the bottom. *4CL*, 4-coumarate:CoA ligase; *CSE*, caffeoyl shikimate esterase; *APX*-*C3H*, ascorbate peroxidase-coumarate 3-hydroxylase; *CCoAOMT*, caffeoyl-CoA *O*-methyl-transferase; *F5H*, ferulate 5-hydroxylase; *GT*, glycosyltransferase; *WND*, wood-associated NAC-domain; *ARK*, ARBORKNOX; *CesA*, cellulose synthase; *KOR*, KORRIGAN; *CSI*, cellulose synthase interacting protein, *GT43*, xylan xylosyltransferase; *GUX*, glucuronyltransferase; *GXM*, glucuronoxylan *O*-methyltransferase; *XOAT*, xylan *O*-acetyltransferase; *GATL*, galacturonosyl-transferase-like; *GH9B*, class B endoglucanase; *CSLA*, mannan synthase; *GMPP*, GDP-mannose pyrophosphorylase; *GAUT*, galacturonosyltransferases; *AAT*, arabinosyltransferase; *BXL*, β-xylosidases/α-arabinofuranosidase; *GH*, glycosyl hydrolase; *MAN*, mannase; *CEL*, cellulase; *XTH*, xyloglucan endotransglucosylase; *ASD*, arabinofuranosidase; *PME(I)*, pectin methyl-esterase (inhibitor); *AGP*, arabinogalactan protein; RNP, ribonucleoprotein.

Transcript levels of many lignin biosynthetic genes, including *4CL1* paralog *4CL5*, were not significantly changed in the *4cl1* mutants. However, the genome duplicates *F5H1* and *F5H2* encoding ferulate/coniferaldehyde 5-hydroxylases were significantly downregulated, and they are the only known monolignol pathway genes besides *4CL1* to show such patterns (Figure 4A). As F5Hs occupy the branch-point into S lignin biosynthesis, their downregulation is consistent with the preferential reduction of S lignin in *4cl1* mutants. Other genes implicated in lignification, including a UDP-glycosyltransferase (Potri.014G096100) orthologous to *Arabidopsis* UGT72B1 involved in monolignol glycosylation (Lin et al., 2016) and a laccase (Potri.010G193100) involved in oxidative coupling of monolignols (Ranocha et al., 2002; Lu et al., 2013), were also downregulated in the mutants (Figure 4A). In contrast, transcript levels of two lignin genes increased significantly, and they encode enzymes acting immediately upstream (caffeoyl shikimate esterase, CSE1) and downstream (caffeoyl-CoA *O*-methyltransferase, CCoAOMT1) of the 4CL reaction with caffeic acid. Both enzymes are encoded by genome duplicates with similar expression levels in poplar xylem and, in both cases, only one copy each was affected in the *4cl1* mutants (Figure 4A). This suggests involvement of CSE1 and CCoAOMT1 in homeostatic regulation of caffeic acid and caffeoyl-CoA in response to *4CL1*-KO. Recently, a cytosolic ascorbate peroxidase (APX) was shown to possess 4-coumarate 3-hydroxylase (C3H) activity and convert 4-coumaric acid to caffeic acid in both monocots and dicots, supporting an alternative route in lignin biosynthesis involving free phenolic acids (Barros et al., 2019). Two orthologs of this dual-function APX-C3H are present in the poplar genome, and one of them (Potri.009G015400, APX-C3H1) was significantly upregulated in the mutants (Figure 4A). Although catalytic properties remain to be confirmed, upregulation of the putative poplar *APX-C3H1* may also contribute to elevated accumulation of caffeic acid and its derivatives.

### Cell wall remodeling and detoxification responses in the *4cl1* mutants

Genes downregulated in the KO mutants showed an overrepresentation of Gene Ontology (GO) terms associated with cell wall biogenesis and polysaccharide biosynthesis (Figure 4C). Examples include transcription factors (TFs), such as top-level WND (wood-associated NAC-domain) proteins, WND1A and WND5A, and downstream TFs, NAC154, NAC156, MYB10, MYB128, MYB167, and MYB221, that regulate lignin, cellulose and xylan biosynthesis (Ye and Zhong, 2015) (Figure 4A). There was widespread downregulation of genes involved in the biosynthesis of all major cell wall glycans, including cellulose, xylans, and pectins (Figure 4A). In contrast, genes encoding cell wall-modifying enzymes or cell wall-loosening proteins were significantly upregulated (Figure 4B). The results are consistent with the altered cell wall polysaccharide composition and extractability, and provide molecular bases for cell wall remodeling as a result of impaired lignification in the mutants.

GO categories associated with detoxification, sulfur assimilation, and oxidation-reduction processes were over-represented among genes upregulated in the mutants (Figure 4C). Of particular interest are genes involved in sulfur assimilation into Cys for synthesis of other sulfur-containing compounds, including glutathione. Upregulation of genes encoding glutathione *S*-transferases (GSTs), redox-active proteins, and enzymes in the glutathione-ascorbate cycle underscores the likelihood of redox adjustments in the mutants (Figure 5). Also increased in the mutants were transcript levels of precursor genes encoding phytosulfokines, disulfated pentapeptides that have been implicated in growth and defense signaling (Sauter, 2015) (Figure 5). Together, these results suggest that cellular redox balance, oxidative stress responses and sulfur homeostasis adjust to changes in thioesterification of the lignin pathway.

**Figure 5.**
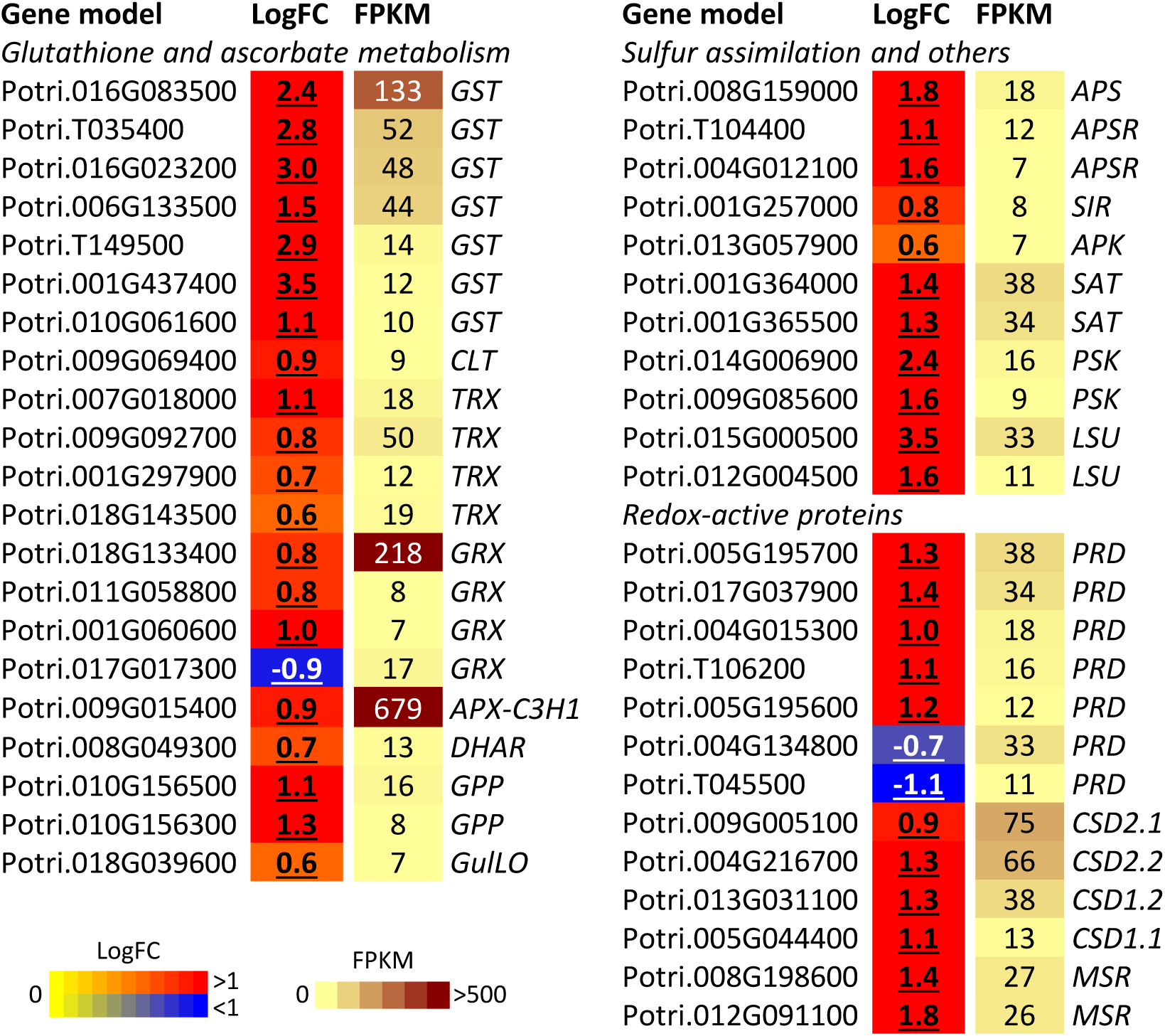
Transcriptional responses of detoxification genes in the mutants. Expression response heatmaps of genes associated with glutathione-ascorbate metabolism, sulfur assimilation and antioxidant systems. Data presentation is the same as Figure 4. Only genes with control FPKM ≥5 (or ≥10 in the case of *GSTs* and *PRDs*) are shown. *GST*, glutathione *S*-transferase; *CLT*, chloroquine-resistance-like (glutathione) transporter; *TRX*, thioredoxin; *GRX*, glutaredoxin; *APX-C3H1*, ascorbate peroxidase-coumarate 3-hydroxylase; *DHAR*, dehydroascorbate reductase; *GPP*, galactose-1-phosphate phosphatase; *GulLO*, gulonolactone oxidase; *APS*, ATP sulfurylase; *APSR*, adenosine-phosphosulfate reductase; *SIR*, sulfite reductase; *APK*, adenosine-phosphosulfate kinase; *SAT*, Ser acetyltransferase; *PSK*, phytosulfokine; *LSU*, response to low sulfur; *PRD*, peroxidase; *CSD*, Cu/Zn superoxide dismutase; *MSR*, Met sulfoxide reductases.

### Conditional involvement of *4CL5* in lignin biosynthesis

The *4cl1* mutants somehow accrue ∼80% of wild-type lignin levels (Table 1), even though *4CL1* transcripts normally comprise >80% of xylem-expressed *4CL* transcript levels in poplar (Supplemental Figure 1). The only other xylem-expressed 4CL isoform, 4CL5, must therefore sustain lignin biosynthesis in the absence of 4CL1. Interestingly, CRISPR-KO of *4CL5* did not change either lignin content or S/G ratio of the stem wood (Supplemental Figure 3), suggesting only a minor or conditional role in lignification under normal growth conditions. As *4CL5* transcript abundance was not significantly altered in the *4cl1* mutants (Figure 4A), other mechanism(s) must exist for *in vivo* enhancement of its function. We constructed sample-specific gene coexpression networks (see Methods) to compare the expression contexts of *4CL1* and *4CL5*, and their shifts in response to *4CL1*-KO (Supplemental Figure 4). The WT network placed *PAL2* and *4CL1* along with *NAC154* and *MYB128* in the brown module, *4CL5* and *APX-C3H1* in the yellow module, and *PAL5, CSE1, CCoAOMT1, F5H1*, and *F5H2* along with *MYB167* and *MYB221* in the turquois module (Figure 6, Dataset 1). In the *4cl1*-KO network, *4CL1, CCoAOMT2, F5H1, F5H2, NAC154, MYB128*, and *MYB167* were assigned to the turquois module, and *PAL2, PAL5, 4CL5, CSE1* and *CCoAOMT1* to the blue module (Figure 6, Dataset 1). The data support distinct regulation of *4CL1* and *4CL5*, and suggest transcriptional rewiring of *4CL5, CSE1*, and *CCoAOMT1* to the same coexpression module as part of a compensatory response to sustain lignification in the *4cl1* mutants.

**Figure 6.**
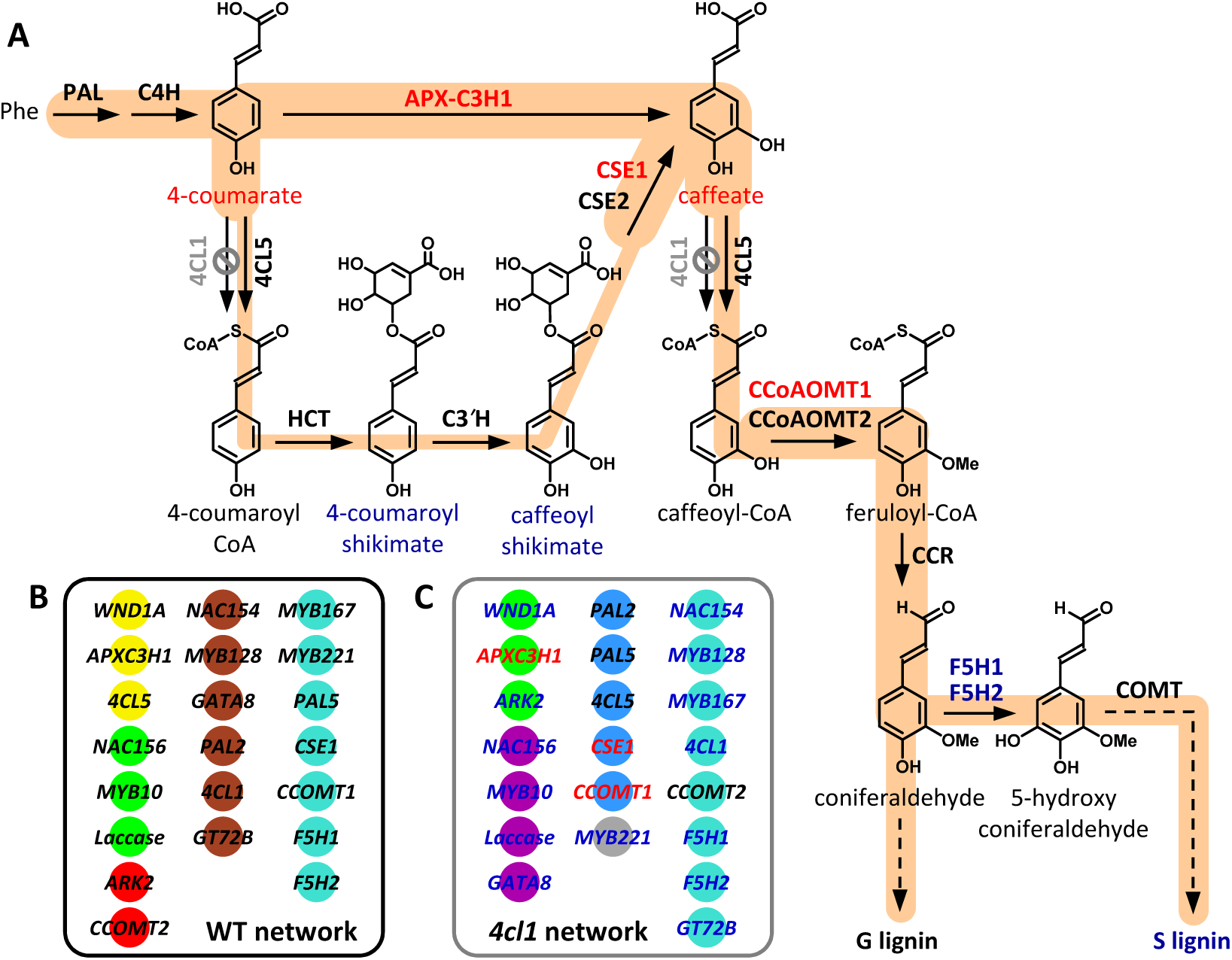
Schematic of the transcriptional and metabolic adjustments of lignin biosynthesis in *4cl1* mutants. **A**, The major monolignol biosynthetic routes are shown with relevant pathway steps and intermediates discussed in this study. Red and blue fonts indicate higher and lower abundances, respectively, in the *4cl1* mutants. The slightly reduced G lignin is shown in black to contrast with the drastically reduced S lignin in blue. Orange background shading depicts the pathway flows, with thickness representing relative transcript and metabolite level changes detected in the mutants. **B-C**, Coexpression module assignments for lignin pathway genes and TFs in control (**B**) and mutant (**C**) networks. Genes are arranged and color-coded by modules they were assigned to, and red and blue fonts in **C** indicate up- or down-regulation in the mutants. Abbreviations are the same as in Figure 4, except for *CCOMT* = *CCoAOMT*.

## DISCUSSION

### *4CL*-KO effects on lignin composition and enzymatic hydrolysis differ between poplar and herbaceous species

The cell wall polymer lignin provides structural integrity and protection to plants (Weng and Chapple, 2010). Its abundance and structural complexity, however, render it a negative factor in forage quality and inhibit its removal in cellulosic biomass utilization for pulping and biofuels production (Boerjan et al., 2003). Studies with wild *Populus trichocarpa* populations (Studer et al., 2011), *brown midrib* sorghum and maize mutants (Saballos et al., 2008; Xiong et al., 2019), *Arabidopsis* T-DNA mutants (Van Acker et al., 2013), and transgenic alfalfa and *Populus* (Chen and Dixon, 2007; Mansfield et al., 2012; Wang et al., 2018) have reported a major, and negative, effect of lignin content on enzymatic sugar release, with lignin S/G ratio playing a minor role. For instance, KO of lignin-associated *4CL* led to reduced lignin content and improved enzymatic saccharification in *Arabidopsis*, maize, sorghum, and switchgrass (Saballos et al., 2008; Van Acker et al., 2013; Park et al., 2017; Xiong et al., 2019). However, for the poplar *4cl1* mutants investigated here, similar degrees of lignin reduction had at best negligible effects on sugar release by multiple pretreatment methods (Figures 2G and 2H). The S/G ratio was reduced by ∼30% in poplar mutants due to drastically reduced S lignin abundance (Table 1), but was increased in *Arabidopsis*, switchgrass, sorghum, and maize mutants owing to strong reductions of G lignin contents (Saballos et al., 2008; Van Acker et al., 2013; Park et al., 2017; Xiong et al., 2019). Poplars and other angiosperm trees with significant secondary growth typically contain S-rich lignin (S/G >1.5), whereas herbaceous species are usually G-rich (e.g., Sun et al., 2012). The biased effects of *4CL*-KO on S and G lignin flux of woody and herbaceous species therefore seem to track with canalized taxonomic determinants of lignin S/G ratio. S lignin is known to be associated with more chemically labile linkages and possess a higher reactivity during pulping as well as various biomass pretreatment regimes (Studer et al., 2011; Mansfield et al., 2012). It thus appears that the potential gain in enzymatic hydrolysis brought about by reduced lignin was offset by decreased S/G ratio in the poplar *4cl1* mutants. We interpret these results to suggest that lignin reduction predominantly in the form of S units does not improve biomass saccharification.

### Caffeic acid homeostasis is sensitive to *4CL1*-KO and modulates 4CL5 compensation

Plasticity in the lignin biosynthetic pathway as evidenced by multiple possible routes (Barros et al., 2019) (Figure 6) may accommodate developmental, environmental, or species-dependent variations in lignification. 4CLs, for instance, can utilize multiple hydroxycinnamate substrates *in vitro*, but how substrate utilization is regulated *in vivo* is largely unknown. 4CL activation of 4-coumarate is obligatory for production of the downstream intermediates, 4-coumaroyl and caffeoyl shikimate esters, in lignin biosynthesis (Schoch et al., 2001; Hoffmann et al., 2004; Coleman et al., 2008), and this was confirmed by greatly depleted shikimate ester levels in the *4cl1* mutants (Table 2). Like many Class I 4CLs involved in lignin biosynthesis, poplar 4CL1 also accepts as substrate caffeic acid, which is the most abundant free hydroxycinnamate in poplar xylem (Harding et al., 2002; Chen et al., 2013) (Table 2). The significant increases of free caffeic acid and its hexose conjugates in the mutant xylem are consistent with a substrate buildup due to impaired 4CL1 conversion. Caffeic acid accrual can also be attributed to conversion from surplus 4-coumaric acid, another 4CL1 substrate, by the upregulated APX-C3H1, as well as from residual caffeoyl shikimate via CSE1. No incorporation of caffeates or caffeyl alcohol into the lignin polymer was evident in the NMR spectra (even at levels nearer the baseplane than those plotted in Figure 1). Besides the reduced S/G ratio and consequent changes in the interunit linkage distribution that were largely as expected, and the elevation of *p*-hydroxybenzoate levels, there were no other evident structural changes in the *4cl1* lignin, and no obviously new components as has often been observed in mutants and transgenics (Ralph and Landucci, 2010).

Several lines of evidence suggest that increased caffeic acid accrual in the *4cl1* mutants was not merely due to precursor buildup, but to a tightly regulated compensatory mechanism to sustain lignification. First, the *4cl1* mutants exhibit ∼20% lignin reduction; that the reduction is not more severe means that substantial levels of lignin biosynthesis must be upheld by 4CL5, the remaining xylem isoform. Second, xylem transcript and protein abundances of 4CL5 are several-fold lower than those of 4CL1 (Wang et al., 2018) (Figure 4A). Third, *4CL1* and *4CL5* exhibited distinct coexpression patterns with other genes and TFs implicated in lignin biosynthesis (Figure 6). These, along with the silent lignin phenotype of *4cl5* mutants during normal growth (Figure S2), all point to 4CL5’s conditional involvement in lignin biosynthesis under specific circumstances. Fourth, all three lignin genes upregulated in the *4cl1* mutants, *CSE1, APX-C3H1* and *CCoAOMT1*, occupy pathway steps immediately upstream and downstream of caffeic acid-to-caffeoyl-CoA activation (Figure 6). This is in sharp contrast to *Arabidopsis 4cl1* mutants in which significant upregulation of early pathway genes *AtPAL2, AtC4H* and *AtC3′H* was reported (Vanholme et al., 2012). Our finding that *4CL5, CSE1*, and *CCoAOMT1* belonged to the same coexpression module in the *4cl1*-specific network highlights a specific involvement of *CSE1* and *CCoAOMT1* in the distinct lignin compositional changes in the poplar *4cl1* mutants. Elevated *CSE1* expression would be expected to direct residual lignin biosynthetic pathway fluxes toward caffeic acid, while also relieving the buildup of potential 4CL inhibitors, 4-coumaroyl and caffeoyl shikimates, in the process (Lin et al., 2015). The increased supply of caffeic acid would be necessary for 4CL5 catalysis to commence because its estimated *K*_m_ value is several-fold higher than that of the predominant 4CL1 (Chen et al., 2013). The observed 3-fold increase in free caffeic acid levels in the *4cl1* mutants is in general agreement with this scenario. Upregulation of the downstream *CCoAOMT1* could further drive the caffeate-to-caffeoyl-CoA flux toward lignification. This, along with caffeic acid glycosylation, could prevent overaccumulation of caffeic acid that may cause substrate self-inhibition of 4CL5 (Chen et al., 2013). Thus, transcriptional co-activation of *CSE1* and *CCoAOMT1* in effect altered the biochemical milieu, enabling the low-affinity 4CL5 to engage in a compensatory function to sustain lignification, even though *4CL5* expression was not increased in the *4cl1* mutants. Regardless, the 4CL5 pathway was not very efficient and could only partially restore lignification in the *4cl1* mutants. We further argue that the 4CL5 compensatory pathway is primarily channeled toward G lignin biosynthesis due to downregulation of *F5H* paralogs. The data thus provide a mechanistic basis for the biased effect on S lignin accrual in the poplar *4cl1* mutants.

### Crosstalk between lignification, redox regulation, and sulfur metabolism

Caffeic acid is among the most potent antioxidants of the plant-derived phenolics, with radical scavenging and metal chelation activities to avert potential damaging effects of reactive oxygen species (Chen and Ho, 1997; Perron and Brumaghim, 2009). Its hyper-accumulation in *4cl1* mutants coincided with upregulation of not only *APX-C3H1* and *CSE1*, but also *GSTs, PRDs, CSDs, MSRs*, and genes encoding enzymes of the glutathione-ascorbate cycle, all of which have been implicated in detoxification of hydrogen peroxide and superoxide (Rouhier et al., 2008; Gao et al., 2010; Molina-Rueda et al., 2013; Rey and Tarrago, 2018; Barros et al., 2019). Coordinated regulation of multiple antioxidant systems may be crucial to maintain redox balance under the presumably highly oxidative cellular environment resulting from disturbed lignin biosynthesis. GSTs also mediate conjugation of glutathione to xenobiotics or electrophilic natural products, including phenylpropanoids, for sequestration and/or transport (Dean et al., 1995; Dixon et al., 2010). Thus, the widespread upregulation of *GSTs* in the *4cl1* mutants could also be linked to detoxification of excess phenylpropanoids.

Besides its roles in redox balance and detoxification, glutathione as an abundant low-molecular-weight thiol metabolite is intimately associated with sulfur homeostasis (Takahashi et al., 2011). Many sulfur assimilation pathway genes can be induced by sulfur starvation, or upon Cd exposure, which triggers synthesis of glutathione and its oligomeric phytochelatins for metal chelation, resulting in sulfur depletion (Scheerer et al., 2010; Honsel et al., 2011). Interestingly, two genes belonging to the plant-specific *LSU* (response to low sulfur) family were significantly upregulated in the mutant xylem (Figure 5), hinting at potential sulfur limitation there. Sulfur assimilation also supplies sulfate donors for sulfation of diverse peptides, proteins, and metabolites (Jez et al., 2016). We observed elevated expression of precursor genes encoding sulfated pentapeptides called phytosulfokines (Figure 5). Sulfation of the tyrosine residues is indispensable for the biological activities of phytosulfokines (Matsubayashi and Sakagami, 1996), which range from growth promotion to defense signaling, including long-distance signaling via xylem sap for stress adaptation (Okamoto et al., 2015). Two unusual sulfate esters, caffeic acid sulfate and ferulic acid sulfate, also increased in the mutants, though with unknown function (Table 2). Taken together, elevated levels of glutathione conjugation, phytosulfokines, and phenolic acid sulfates might have depleted available pools of reduced sulfur in the *4cl1* xylem, which in turn activated the sulfur assimilation pathway.

## CONCLUSIONs

Our data support differences in regulation and perturbation of lignin biosynthesis between *Populus* and herbaceous species. We identified the juncture between caffeic acid and caffeoyl-CoA as being highly sensitive to *4CL1*-KO, while also key to modulating the compensatory function of 4CL5. Transcriptional, metabolic, and biochemical coordination of the compensatory pathway likely underscores the species-specific lignin perturbation response reported here. The work raises the enticing possibility that such lineage-specific capability might have originated from taxon-dependent developmental plasticity that warrants future investigation.

## MATHERIALS AND METHODS

### Plant materials

The *4cl1* mutants and Cas9-only vector controls were reported previously (Zhou et al., 2015). The frameshift mutations in the gRNA target site of *4CL1* (GAGGATGaTaAAaTCTGGAG<u>GGG</u>, the protospacer adjacent motif, PAM, is underlined and sequence polymorphisms with *4CL5* in lowercase) have been reconfirmed using independently collected leaf samples (Bewg et al., 2018). *4CL5*-KO plants were produced by targeting the homologous site (GAGGATGtTgAAgTCTGGA<u>GGG</u>). The construct was prepared according to Jacobs and Martin (2016), except two oligos tailed with *Medicago truncatula* U6 (AACTCCAGACTTCAACATCCT<u>CAAGCCTACTGGTTCGCTTGA</u>) or scaffold sequence (GAGGATGTTGAAGTCTGGA<u>GTTTTAGAGCTAGAAATAGCAAGTT</u>, underlined) were used in a half (10 μL) reaction containing 0.005 pmol of linearized p201N vector and 0.1 pmol of inserts (U6, scaffold and the two gRNA oligos) using the Gibson Assembly Master Mix (NEB). *Agrobacterium*-mediated transformation of *P. tremula* x *alba* INRA 717-1B4 was performed according to Leple et al. (1992). Editing patterns were determined by amplicon-sequencing using *4CL1*/*4CL5* consensus primers (<u>GTTCAGACGTGTGCTCTTCCGATC</u>TAGCACCGGTTGTHCCA and <u>CCTACACGACGCTCTTCCGATCT</u>GAGGAAACTTRGCTCTGAC) tailed with Illumina sequencing primers (underlined) to check for both on-target and off-target cleavage as described (Zhou et al., 2015). Indexed samples were pooled and sequenced as a spike-in on an Illumina NextSeq500 at the Georgia Genomics and Bioinformatics Core, University of Georgia, and the data were analyzed by AGEseq (Xue and Tsai, 2015). Developing xylem scrapings were collected from ∼2 m trees at 15-30 cm from the top and snap-frozen in liquid nitrogen. The rest of the stem was debarked and air-dried for wood chemical and physical analyses. Plants were vegetatively propagated by stem cuttings for use in histological analysis.

### Histology

Stem cross sections (100 μm) were prepared from young shoots using a Vibratome (Ted Pella). Sections were stained with 0.05% (w/v) toluidine blue or 2% (w/v) phloroglucinol in acid ethanol (1 M HCl:95% ethanol = 1:1, v/v), and images were acquired using a Zeiss Axioskop-50 microscope equipped with a Leica DFC500 digital camera.

### Wood specific gravity and acoustic velocity

Three ∼8 mm longitudinal stem wood sections were obtained from the base of each sample for nine control (three WT and six Cas9 lines) and 12 mutant samples. Specific gravity (wood density divided by the density of water) was measured by water displacement after the sections were saturated with water (ASTM International, 2017). Acoustic velocity was measured on the samples prior to water saturation using a time of flight instrument (SoniSys) equipped with two 1-MHz transducers to send and receive the acoustic signal. The specific gravity and acoustic velocity values from the three sections were averaged for each sample, and statistical differences between groups were determined by Student’s *t* test.

### Wood chemistry, saccharification, and NMR analysis

The wood samples were ground to pass a 40-mesh screen using a Wiley mill, followed by 99 cycles of ethanol extraction using a Soxhlet extraction unit (Buchi) and air dried. Extractive-free wood meal was used for Klason lignin (Swamy et al., 2015) and crystalline cellulose content determination according to Updegraff (1969). Lignin composition was determined by thioacidolysis (Foster et al., 2010), matrix polysaccharide composition by trifluoroacetic acid hydrolysis (Foster et al., 2010), and glycosyl composition by methanolysis followed by trimethylsilylation (Harding et al., 2018). High-throughput saccharification assays were performed using the BESC method with hot water pretreatment at 180 °C for 40 min (Selig et al., 2010), as well as the GLBRC methods using hot water, dilute acid (2% H_2_SO_4_) or dilute base (6.25 mM NaOH) pretreatment at 90 °C for 3 h (Santoro et al., 2010). NMR analysis using ball-milled whole cell wall residues or enzyme lignin following cellulase digestion was performed as detailed (Kim and Ralph, 2010; Mansfield et al., 2012). Unless otherwise noted, these analyses were performed for five controls (three WT and two vector controls) and five *4cl1* mutants.

### Glycome profiling

Extractive-free wood meals were sequentially extracted with 50 mM ammonium oxalate, 50 mM sodium carbonate, 1 M KOH, 4 M KOH, acid chlorite and 4 M KOH to obtain cell wall fractions for screening with a panel of plant glycan-directed monoclonal antibodies (mAbs) (Pattathil et al., 2010), as described (Pattathil et al., 2012). Statistical analysis of mutant effects was performed with Limma (Smyth, 2005) after excluding mAbs with hybridization intensities <0.1 in all samples (three WT, three vector controls and six *4cl1* mutants). mAbs that showed significantly different signal intensities between mutant and control samples (*P* ≤0.01 and fold-change ≥1.5) were visualized by heatmaps using *R*.

### Phenolic profiling

Developing xylem scrapings were ground to a fine powder under liquid nitrogen, aliquoted and stored at −80 °C. One aliquot per sample was freeze-dried and 10 mg (dry weight) of tissue were extracted with 400 μL chloroform:methanol (1:1, v/v) in an ultrasonic bath with pre-ice-chilled water for 30 min, followed by addition of 200 μL water, vortexing and centrifugation for phase separation. The aqueous phase was transferred to a new tube, and an aliquot of 100 μL was evaporated to dryness in a Centrivap (Labconco) for shipment to the VIB. The residues were resuspended in 200 μL of cyclohexane:water (1:1, v/v) and 15 μL were injected onto an Acquity UHPLC equipped with a Synapt Q-TOF (Waters) for chromatographic separation and mass spectrometric detection of phenolic metabolites following conditions and settings detailed previously (Saleme et al., 2017). A total of 10 control (three WT and seven Cas9) and 14 *4cl1* samples were analyzed. Significant differences between the two groups were declared for peaks that had average signal intensities ≥1000 counts in at least one group, with *P* ≤0.01 and fold-change ≥2. Compound annotation was based on matching *m/z*, retention time and MS/MS fragmentation (Supplemental Figure 5) against an in-house library as well as literature data.

### RNA-seq analysis and sample-specific coexpression network construction

RNA was extracted from frozen xylem powder using the Direct-zol RNA Miniprep kit (Zymo) with PureLink Plant RNA Reagent (Life Technologies) for Illumina RNA-Seq library preparation and NextSeq 500 sequencing as described (Harding et al., 2018). Six control (three WT and three Cas9) and six *4cl1* mutant samples were sequenced to obtain 8.3-12.1 million (M) paired-end 75-bp reads per sample, except vector control Cas9-39 for which >97M reads were obtained and only randomly sampled 16M reads were used for data analysis. After preprocessing for quality control, reads were mapped to the variant-substituted *Populus tremula* × *alba* genome v1.1 as detailed previously (Xue et al., 2015). Genes satisfying the criteria of FPKM ≥5 in at least four out of six replicates of at least one group, average FPKM ≥10 in at least one group, *Q* ≤0.05 and fold-change ≥1.5 were used for GO enrichment analysis by topGO (Alexa and Rahnenfuhrer, 2010) with Fisher’s exact test, and the negative log_10_ transformed *P* values were visualized as heatmaps. For coexpression network analysis, relaxed thresholds (*P* ≤0.05, FPKM ≥5) were used to obtain 5512 genes that differed between control and *4cl1* plants. The average expression values of the 5512 genes were calculated for the control and *4cl1* mutant groups. We then adapted the approach of Liu et al. (2016) for construction of sample-specific networks. Briefly, a group of reference samples was assembled from our previous studies unrelated to lignin pathway gene manipulation (Swamy et al., 2015; Xue et al., 2016). Only unstressed xylem samples were included (n=35). Expression values of the 5512 genes were extracted from the 35 samples, and the averaged expression values of control or *4cl1* mutant samples were added separately to create two datasets (n=36 each) for construction of the control-and mutant-specific networks. For simplicity, we refer to the former containing the control group (WT and Cas9 lines) as the WT network. Pairwise Spearman correlation coefficients of both datasets followed a normal distribution (Supplemental Figure 4) and were used for weighted gene coexpression network analysis using the WGCNA *R* package (Langfelder and Horvath, 2008) as detailed in Xue et al. (2016), with a soft threshold of 10. The coexpression relationships were assessed by hierarchical clustering using the topological overlap measure and modules were determined with a dynamic tree cutting algorithm (Supplemental Figure 4). The list of 5512 genes, their expression differences between control and *4cl1* mutants, and sample-specific network module assignments are provided in Dataset 1.

## ACCESSION NUMBERS

The RNA-seq data has been deposited to the NCBI Sequence Read Archive under accession PRJNA589632.

## ACKNOWLEDGMENTS

The authors thank Gilles Pilate of INRA, France for providing poplar clone INRA 717-1B4, Yenfei Wang for assistance with transgenic plant production, Eli McKinney for greenhouse plant care, Nicholas Rohr and Bob Schmitz for Illumina RNA library construction, Jacob Reeves for assistance with data processing and SRA submission, the Georgia Genomics and Bioinformatics Core for Illumina NextSeq sequencing, the Complex Carbohydrate Research Center Analytical Services and Training Laboratory for pyMBMS analysis, and Cliff Foster of GLBRC for cell wall analyses.

## SUPPLEMENTAL DATA

**Supplemental Figure 1.** Class I 4CL gene family members in *Populus*.

**Supplemental Figure 2.** Glycome profiling analysis of all six cell wall fractions

**Supplemental Figure 3.** Characterization of *4cl5* knockout mutants

**Supplemental Figure 4.** Sample-specific coexpression network analysis

**Supplemental Figure 5.** MS/MS spectra of metabolites listed in Table 2

**Supplemental Dataset 1.** List of differentially expressed genes and their network module assignments

